# A potential *Toxoplasma gondii* lipoxygenase is necessary for virulence and associated with the host immune response

**DOI:** 10.1101/2022.05.12.491751

**Authors:** Carlos J. Ramírez-Flores, Andrés M. Tibabuzo Perdomo, Sarah K. Wilson, Carolina Mendoza Cavazos, Billy Joel Erazo Flores, Katie L. Barnes, Laura J. Knoll

**Author notes:** Address correspondence to: Laura J. Knoll, 3303 Microbial Sciences Building 1550 Linden Drive Madison, WI 53706. All the authors contributed equally to this manuscript and share first authorship (Carlos J. Ramírez-Flores, Andrés M. Tibabuzo Perdomo, Sarah K. Wilson).

## Abstract

While the asexual cycle of *Toxoplasma gondii* can occur in any warm-blooded animal, the sexual cycle is restricted to the feline intestine. We previously determined that because cats lack delta-6-desaturase activity in their intestines, they build up excess linoleic acid, which signals *T. gondii* to undergo sexual development. We hypothesized that *T. gondii* oxygenates linoleic acid to signal sexual development, so we examined the *T. gondii* genome for potential lipoxygenases (TgLOX) enzymes. We identified seven potential TgLOXs that were at least 100-fold more abundant in the cat intestinal versus the tissue culture tachyzoite stage. Parasites deleted in TgLOX1 (TgΔLOX1) had no significant growth differences in tissue culture fibroblast cells. Because the sexual development assay begins with brain cysts, we infected mice with TgΔLOX1 and were surprised to find that TgΔLOX1 had reduced virulence. The TgΔLOX1 parasitemia was reduced by 3 days postinfection and largely cleared by 7 days postinfection. At 3 days postinfection, the cytokines IFNγ, IL-6, MCP-1, and TNF-α were significantly reduced in TgΔLOX1-infected mice, which prompted us to examine TgΔLOX1 in IFNγKO mice. We found that IFNγKO mice infected with TgΔLOX1 succumbed to acute infection with the same kinetics as the parental and complemented strains, suggesting the role of TgLOX1 in mice was IFNγ dependent. In tissue culture fibroblasts, TgLOX1 was localized within the parasite, but in leukocytes from infected mice and activated macrophages, TgLOX1 was localized in vesicular structures in the host cytoplasm. These results suggest that TgLOX1 in these vesicular structures modifies the host immune response.

**Importance:** Lipoxygenases are enzymes that catalyze the dioxygenation of polyunsaturated fatty acids such as linoleic and arachidonic acid. These modifications create signaling molecules that are best characterized for modulating the immune response. Deletion of the first lipoxygenase characterized for *Toxoplasma gondii* (TgLOX1) generated a less virulent strain and infected mice showed a decreased immune response. This virulence defect was dependent on the mouse cytokine IFNγ. TgLOX1 changes location from inside the parasite in tissue culture conditions to vesicular structures within the host immune cells during mouse infection. These results suggest that TgLOX1 plays a role in the modification of the host immune response in mice.

## Introduction

*Toxoplasma gondii* is one of the most successful parasites in the world, with around 30% of the world’s human population infected (1). Humans are infected congenitally, or by consuming tissue cysts or oocysts. The tissue cysts form during the asexual cycle in the muscle and brain of infected animals, whereas oocysts are formed during the sexual cycle in felines and are shed in the feces. In healthy human hosts, the infection can be asymptomatic due to the rapid immune response, allowing the parasite to pivot to a chronic infection. However, in immunocompromised individuals, the infection could lead to chorioretinitis, blindness, encephalitis, and even death. During the early stages of infection, chemokines are released by infected enterocytes. This chemokine signaling results in leukocyte recruitment, followed by cytokine production and parasite clearance (2). IL-12 plays an important role in the stimulation of interferon gamma (IFNγ) from the natural killer and T cells in *T. gondii* infected mice (3). IFNγ is considered the main mediator of the acute and chronic defense during *T. gondii* infection (4).

The role of oxygenases in controlling immune response has been well studied (5). The main enzymes involved in the production of these signaling molecules are cyclooxygenases, cytochrome P450, and lipoxygenases. These enzymes belong to the oxidase family and their main function is to transfer molecular oxygen to their substrate (6). Each one plays a central role in lipid metabolism, specifically, the catalysis of polyunsaturated fatty acids that lead to the production of prostaglandins, thromboxanes, leukotrienes, and lipoxins (7).

The effect of host lipoxygenases and cyclooxygenases on *T. gondii* has been examined. It has been shown that inhibition of the host COX-2 enzyme resulted in the decrease of parasite burden in the brain and intraperitoneal macrophages of *Calomys callosus* rodents (8). Similar results were observed in human trophoblasts and villous explants treated with COX-2 inhibitors (9). Host lipoxygenases, appear to be involved in the production of anti-inflammatory responses. When mice are exposed to *T. gondii*, the production of a specialized pro-resolving mediator, Lipoxin A_4_ (LXA_4_), increases. Mice deficient in 5-lipoxygenase, which synthesizes LXA_4_, die at the beginning of chronic infection due to uncontrolled inflammation (10, 11). These studies show the importance of lipid mediators in the control of *T. gondii* infection.

The role of oxygenases encoded by the *T. gondii* genome has not been examined. The pathogenic fungus *Aspergillus flavus* encodes a lipoxygenase (LOX) that is essential for sexual development, suggesting that a LOX-derived metabolite activates the pathway (12). *A. flavus* uses linoleic acid (18:2), but not oleic acid (18:1) to signal sexual development. These results are similar to our finding that *T. gondii* uses linoleic, but not oleic acid to signal sexual development (13). In the present study, we focused on the functional characterization of the novel *T. gondii* lipoxygenase TGME49_315970, hereinafter called TgLOX1. In this work, we describe the unexpected role of TgLOX1 during mouse infection, the cytokine response from wild-type and IFNγ KO mice, as well as the localization of LOX1 within the parasite and immune cells.

## Results

### TgLOX1 is a lipoxygenase expressed in *Toxoplasma gondii*

To study *T. gondii* lipoxygenases potentially involved with sexual development, we performed a bioinformatic screen (Fig 1.). To consider a protein as a lipoxygenase of interest we set two parameters. The C-terminus of the protein must be an Isoleucine or a Valine residue (14, 15). The transcript for the lipoxygenase must be 100-fold upregulated during *T. gondii* sexual stages. Our bioinformatic analysis showed that from the 8322 TGME49 annotated proteins from ToxoDB, 35 had a C-terminal Isoleucine and 260 had a C-terminal Valine residue, a defining characteristic of lipoxygenases. Of the 35 Isoleucine candidates, 7 were at least 100-fold more abundant in the sexual stage comparing it to the tachyzoite stage. From the 260 Valine candidates,11 sequences met the same criteria. With the pool of potential candidates greatly reduced (Table 1), TGME_315970 was selected as our protein of interest due to the high abundance of transcripts (1930 TPM) present in the sexual stage of *T. gondii*.

**Figure 1.**
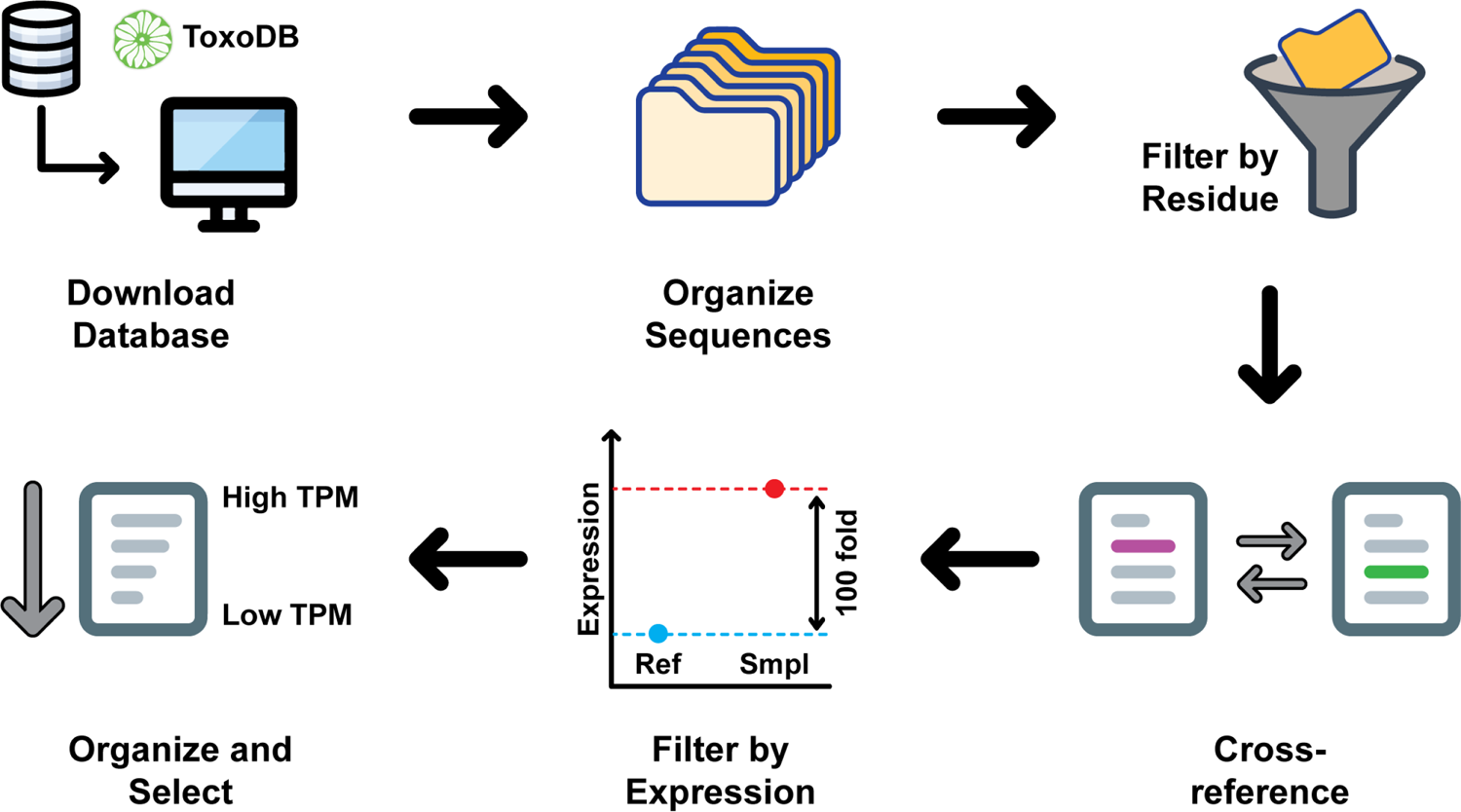
Bioinformatic screening of *T. gondii* proteins to determine potential lipoxygenases. Data were downloaded from ToxoDB and filtered for the presence of an isoleucine or a valine residue in the C-terminus of the protein. An additional sorting was developed by the selection of 100-fold upregulated proteins during *T. gondii* sexual stage compared to the tachyzoite stage. TGME49_315970 was selected as the lipoxygenase of interest due to the high abundance of transcripts during the sexual stage of *T. gondii*.

**Table 1.** Lipoxygenase candidates

| Gene ID | Protein Length | Molecular Weight | C-term residue | Sexual stage (EES5) unique transcripts (TPM) | Tachyzoite stage unique transcripts (TPM) | Fold Change |
| --- | --- | --- | --- | --- | --- | --- |
| TGME49_315970 | 655 | 69105 | I | 1930 | 7.67 | 252 |
| TGME49_218215 | 584 | 64754 | I | 972 | 0.28 | 3470 |
| TGME49_233360 | 174 | 18324 | V | 668 | 5.77 | 116 |
| TGME49_232170 | 745 | 80968 | V | 591 | 0.89 | 664 |
| TGME49_320290 | 534 | 59292 | V | 424 | 0.11 | 3850 |
| TGME49_210400 | 1354 | 144282 | V | 204 | 0.05 | 4080 |
| TGME49_211360 | 863 | 96500 | I | 133 | 0.06 | 2210 |

According to the *T. gondii* transcriptome in feline enteroepithelial stages previously reported (16), the TgLOX1 (TGME49_315970) transcript is 250-fold more abundant during the enteroepithelial stages compared to the tissue culture tachyzoite stage. Given the potential role of TgLOX1 in triggering sexual development, we generated a TgLOX1 depleted strain in PruΔHPT:luciferase parasites. Deletion of the LOX1 gene was confirmed by PCR (Fig. S1, primers in Table S1). We also generated complement strains, both untagged and HA-tagged, by random insertion of the LOX1 gene into the ΔTgLOX1 parasites.

### ΔTgLOX1 parasites show no distinctive phenotype in tissue culture

The parental PruΔHPT:luciferase, ΔTgLOX1, or complement untagged parasite were characterized in tissue culture. We saw no replication differences between the parental and ΔTgLOX1 strains, but slight but significant differences between the ΔTgLOX1 and complement strain at 24 and 36 hours. This small difference could be explained by the random insertion of the TgLOX1 gene in the complemented strain (Fig. S2). The plaque assays did not show significant changes in plaque size or numbers in HFF monolayers when infected with ΔTgLOX1 compared to infection with parental strain but again there was a slight but significant difference between the ΔTgLOX1 and complement strain (Fig S2).

### TgLOX1 is localized within the parasite in HFF cells

To localize TgLOX1, we generated ΔTgLOX1 complement strains with either an N- or C-terminal HA-tag on TgLOX1. We opted for tagging both termini to avoid disturbing the LOX activity given that deletion of the C-terminal isoleucine could alter the lipoxygenase activity of the enzyme (17). We detected TgLOX1-HA by western blot (Fig. 2A), so we localized TgLOX1 protein in intracellular parasites cultured in HFF cells. At 24 hours postinfection, TgLOX1 with either a C or N-terminal HA-tag showed an intracellular localization that did not colocalize with the microneme protein MIC2 (Fig. 2C). Although diffuse labeling was shown while using epifluorescence microscopy, clearer spots within the parasite were detected using a confocal microscope, suggesting a restricted localization in the cytoplasm (Fig. 2B and Movie 1 and 2). The HA-tagged TgPL1 protein (18) was used as a positive control for the HA antibody. It is worth noting that we detected the same distribution of TgLOX1-HA in both C- and N-terminus parasites; however, the abundance of the C-terminal tagged protein was less (Fig 2A and B), perhaps due to the HA-tag interference with the catalytic core and destabilization of the protein (Fig 2B).

**Figure 2.**
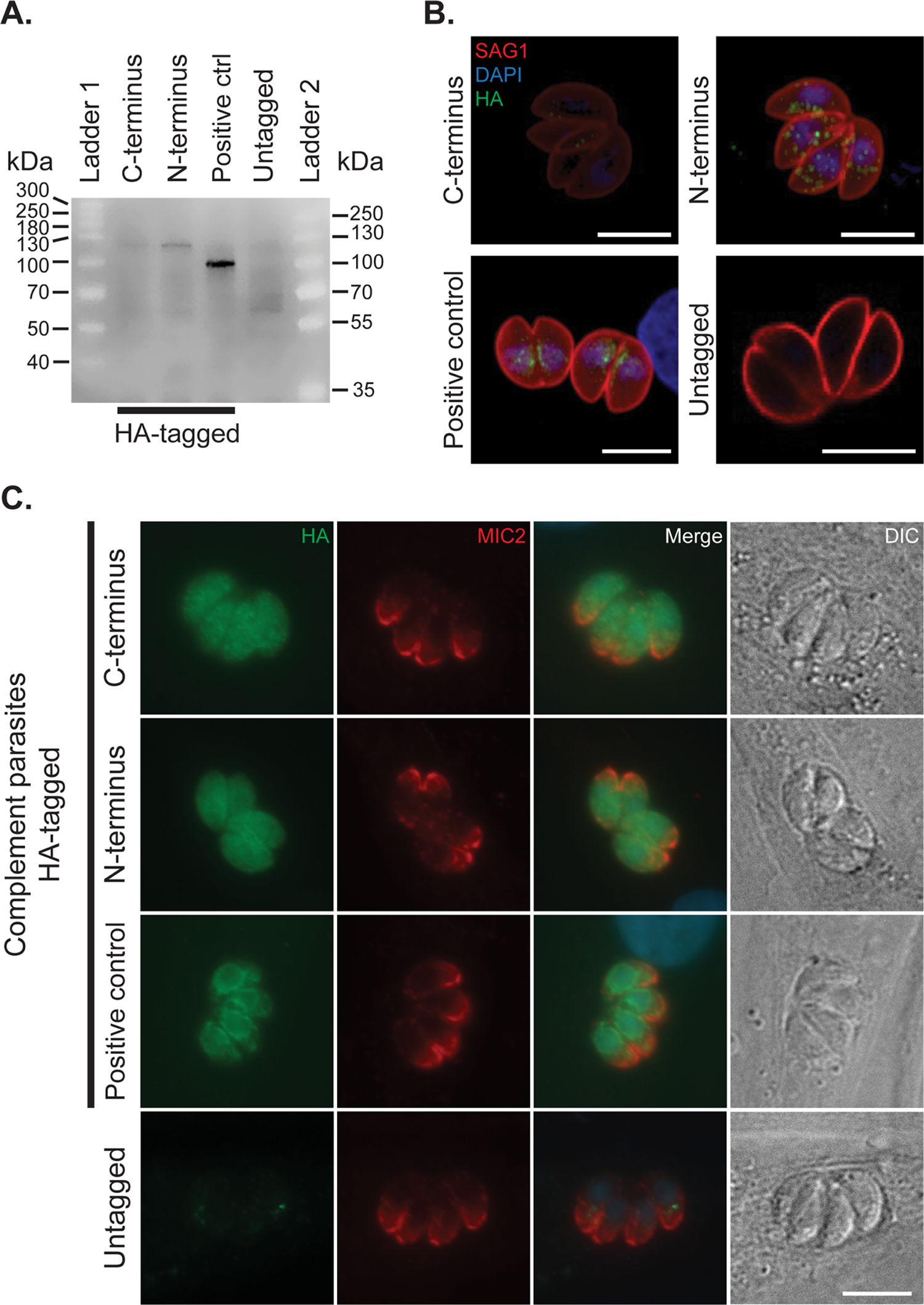
ΔTgLOX1 is expressed and localized in the cytoplasm of intracellular tachyzoites. (A) Parasites expressing HA-tag in the C-terminus or N-terminus were assessed for protein expression by Western analysis with anti-HA antibody. To confirm the molecular heights, two protein ladders were used in the same blot: ladder 1 (ThermoFisher Scientific, Cat. 26625) and ladder 2 (Cat. 26619). TgPL1-HA was a positive control and untagged parasites as a negative control. (B) 3D projections of the cytoplasmic distribution of ΔTgLOX1 (green) and the parasite surface localization of SAG1 (red) in intracellular tachyzoites at 24 hours postinfection and visualized by confocal microscopy. TgPL1-HA (green) was a positive control. (C) Colocalization of MIC2 (red) and ΔTgLOX1 (green) in intracellular tachyzoites at 24 hours postinfection and visualized by epifluorescence microscopy with Differential Interference Contrast (DIC). TgPL1-HA was a positive control (green) and untagged parasites as a negative control. Images for panel C were taken under the same magnification. All of the scale bars are equal to 5 μm.

### ΔTgLOX1 parasitemia is reduced by chronic infection

To evaluate the role of TgLOX1 in the sexual cycle, we needed to obtain mouse brain cysts to initiate the development assay (13). Because all strains contained the luciferase gene, we decided to estimate the chronic infection parasitemia using luciferase activity. NMRI mice were infected intraperitoneally (i.p.) with 1×10^4^ parasites and parasitemia measured at 28 days postinfection. According to the luminescence assay, parasites were found in the brains of the infected mice with parental and untagged complement strains, but no parasites were found in ΔTgLOX1-infected mice (Fig S3A). We then infected NMRI mice with 1×10^5^ parasites, but mice infected with parental and complement parasites did not survive to chronic infection (Fig. S3B). At 28 days postinfection, the brains of ΔTgLOX1-infected mice were stained for *Dolichos Biflorus* Agglutinin (DBA) to visualize the *T. gondii* cyst wall. Almost no cysts were seen in the ΔTgLOX1-infected mice, but the cysts that were seen had intact walls (Fig. S3C). We then infected Swiss Webster mice with 1×10^4^ parasites and monitored for parasitemia at 28 days postinfection; while there was a low luciferase signal in the ΔTgLOX1-infected mice, it was significantly lower than the parental and complement parasites (Fig. S3D).

### ΔTgLOX1 parasites are cleared in mice by day 7 postinfection

To evaluate the role of TgLOX1 during acute infection of *T. gondii,* we infected C57BL/6 mice i.p. with 1×10^5^ parasites and monitored their health over time. Mice infected with parental or untagged complement strains showed signs of declining health by 7 days postinfection, with most becoming moribund (Fig. 3A). Mice infected with the ΔTgLOX1 strain did not show any symptoms of infection by 28 days postinfection. Interestingly, while all of the males needed to be sacrificed, 20% of C57BL/6 female mice recovered and survived out to 28 days postinfection (Fig. 3A).

**Figure 3.**
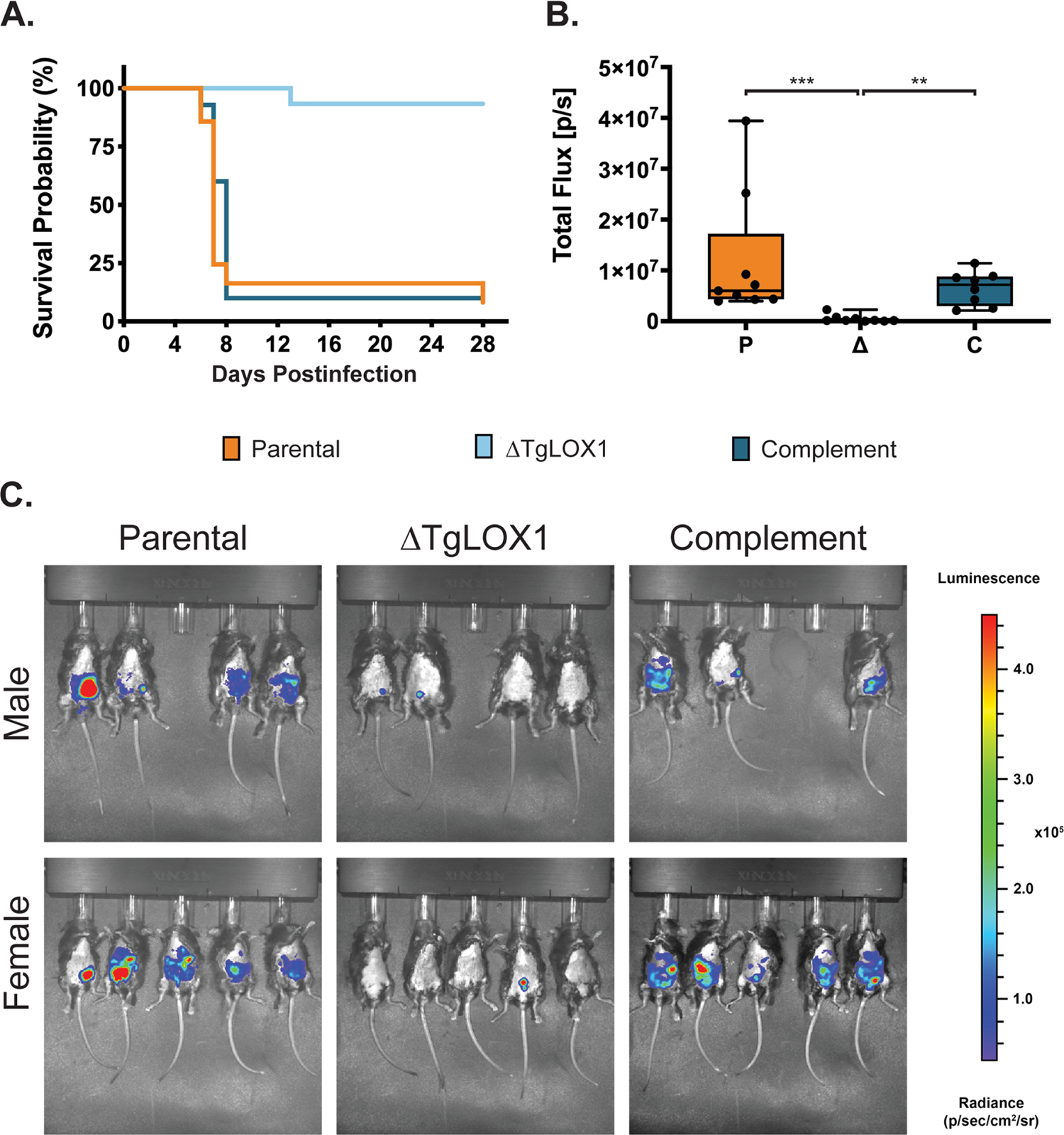
ΔTgLOX1 parasites are avirulent in wild type mice. (A) Shown a combination of two independent experiments of 3-5 C57BL/6-WT mice, either all males or all females, with a total of 6-9 mice per strain. Mice were i.p. infected with 1×10^5^ of each strain and health monitored up to 28 days postinfection. (B and C) Females and males C57BL/6-WT were infected i.p. with 1×10^4^ parasites of each strain and 7 days postinfection they were imaged ventrally by IVIS. (B) Shown the total flux was obtained by measuring the luminescence intensity in the peritoneal cavity of mice. (C) Shown are the images for each strain and separately for gender. For all mice, the abdominal hair was removed to avoid signal interference and the exposure time was the same.

To track the progression of infection in mice, we infected C57BL/6-WT male and female mice with 1×10^4^ parasites of our three strains and proceeded to image them by an *in vivo* imaging system (IVIS) at 7 days postinfection. Mice infected with ΔTgLOX1 parasites showed no or poor signal restricted to the site of the inoculation; in contrast, mice infected with parental and complement strains showed higher levels of parasitemia along the peritoneal cavity (Fig. 3B and C). No sex-dependent differences in parasitemia were found between the male and female mice. The clearing of the ΔTgLOX1 parasites by day 7 postinfection explains the lack of symptoms during acute infection and cysts by day 28 postinfection (Fig. S3).

### Mice infected with ΔTgLOX1 parasites show a reduced cytokine response

The pronounced phenotype that was observed *in vivo* prompted us to investigate the host immune response mounted against the infection. We infected C57BL/6-WT mice with 1×10^4^ parasites of either parental, ΔTgLOX1, or untagged complement strains and collected sera from tail bleeds: before infection, 3 days postinfection, and a final bleed at 7 days postinfection. Sera were then used to quantify the concentration of inflammatory cytokines (IFNγ, IL-6, MCP-1, IL-10, IL-12p70, and TNF-α). Similar to the progression of the infection observed by IVIS at day 7 postinfection, the cytokine response was higher in those mice infected with parental and complement parasites, with no differences between those two groups (Fig. 4). The immune response in ΔTgLOX1-infected mice was significantly reduced compared to the parental and complement strains in all the cytokines tested (Fig. 4). IFNγ showed the highest concentration levels of the four cytokines present in the samples, while other cytokines such as IL-10 and IL-12p70 were not detected. With an infection of 1×10^4^ parasites, the cytokines at 3 days postinfection were in low abundance and were not able to be detected (data not shown).

**Figure 4.**
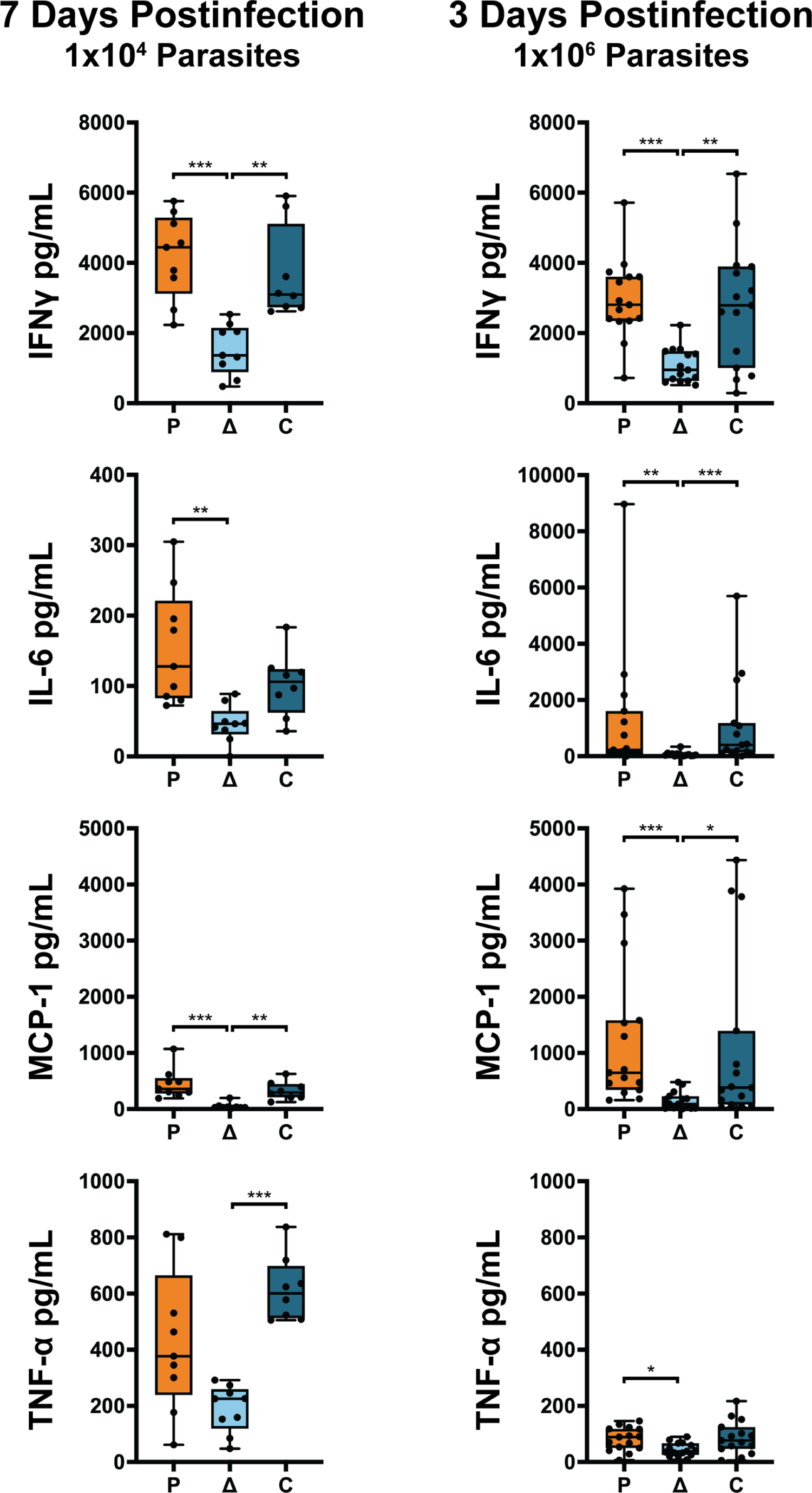
ΔTgLOX1 parasites produced a reduced cytokine response in wild type mice. Blood serum of female and male C57BL/6-WT mice was collected to analyze the cytokine response. Mice were infected with 1×10^4^ or 1×10^6^ parasites of each strain and analyzed at 7 or 3 days postinfection, respectively. The statistics were performed using one way ANOVA in the GraphPad prism software. The *p* value was considered as follows: * <0.05, ** <0.005 and *** <0.0005.

To determine the cytokine response during the early infection, we challenged the mice with a larger dose (1×10^6^ parasites) of each strain and measured the cytokines at 3 days postinfection. As expected from a higher parasite load, the levels of cytokines in the sera increased proportionally. At 3 days postinfection, all the cytokine levels of IL-6, MCP-1, TNF-α, and IFNγ in ΔTgLOX1-infected mice were lower compared to the cytokine level in parental-infected mice (Fig. 4). These results are consistent with the IVIS data, showing that even with this large dose, some mice were able to clear the infection when infected with ΔTgLOX1 parasites, by 3 days postinfection (Fig. S4). However, the avirulent phenotype of ΔTgLOX1 is dosage dependent as mice infected with 1×10^8^ ΔTgLOX1 parasites were symptomatic and moribund by 11 days postinfection (data not shown). These results suggest that TgLOX1 can modulate immune response as early as 3 days postinfection.

### IFNγKO mice are susceptible to ΔTgLOX1 parasites

The IFNγ response is critical for the host defense against *T. gondii* (4). IFNγ levels were significantly lower when mice were infected with ΔTgLOX1 (Fig. 4), so we tested if the immune response modulation by TgLOX1 was IFNγ dependent by infecting IFNγKO mice. Male and female IFNγKO mice were infected with 1×10^4^ parasites of parental, ΔTgLOX1, and complement parasites. All infected mice needed to be sacrificed on days 9 – 10 (Fig. 5A), making it impossible to obtain cysts in their brains, which requires an incubation period of at least 21 days. We repeated the infection with a lower parasite burden of 1000 parasites and then with 100 parasites. In both cases, the spread of the infection was such that all mice died by day 10 postinfection (data not shown). At 7 days postinfection, all of the IFNγKO mice infected with 1×10^3^ parental, ΔTgLOX1, or complement strains showed similar parasitemia (Fig. 5B and C). These results suggest that the ΔTgLOX1 parasites can invade and generate the acute infection efficiently in the absence of IFNγ.

**Figure 5.**
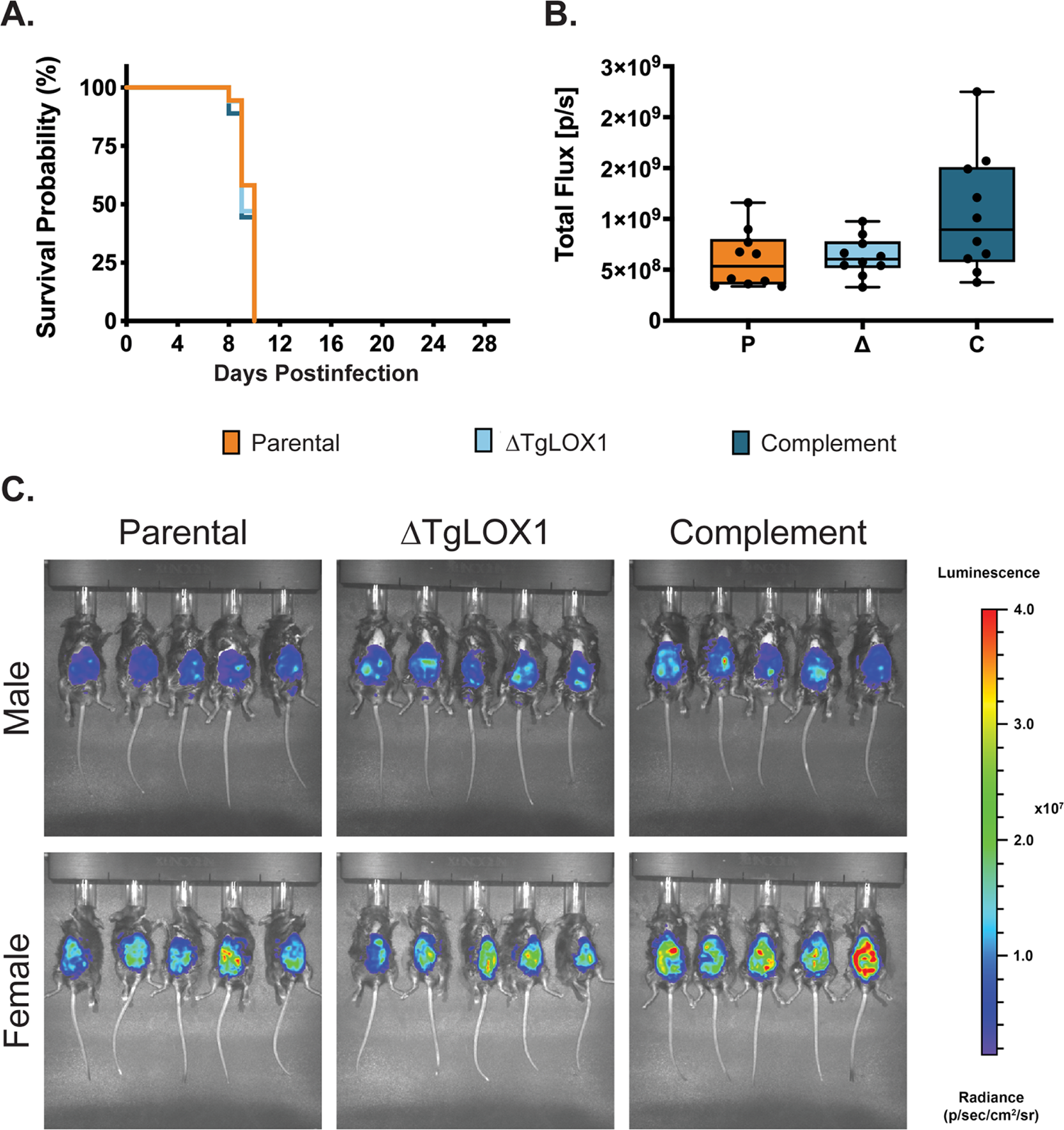
ΔTgLOX1 parasites are virulent in IFNγKO mice. (A) Shown is a combination of two independent experiments of 2-5 IFNγKO mice, either all males or all females, with a total of 7 mice per strain. Mice were i.p. infected with 1×10^4^ of each strain and their health was monitored until they were moribund. (B and C) Females and males IFNγKO were infected i.p. with 1×10^3^ luciferase-expressing parasites of each strain and 7 days postinfection they were imaged ventrally by IVIS. (B) Shown the total flux was obtained by measuring the luminescence intensity in the peritoneal cavity of mice. (C) Shown are the images for each strain and separately for gender. For all mice, the abdominal hair was removed to avoid signal interference and the exposure time was the same.

We analyzed the cytokine response in IFNγKO mice at 7 days postinfection. The results showed that in the absence of IFNγ the concentration of MCP-1 increased ∼10-fold, and IL-6 increased ∼100-fold when compared to the C57BL/6-WT mice, suggesting that other cytokines are upregulated to try to control parasitemia (Fig. S5). It is interesting that TNF-α is significantly lower in ΔTgLOX1-infected mice compared to parental- and complement-infected WT mice, but TNF-α is significantly higher in ΔTgLOX1-infected mice compared to parental- and complement-infected IFNγKO mice. These differences are clearly not protecting IFNγKO mice against *T. gondii* infection as the mice succumb to infection with the same kinetics regardless of which strain they were infected with.

### TgLOX1 is localized in the cytoplasm of infected leukocytes

During pathogen infections in mammals, it is well known that lipoxygenases and their products play an important role in modifying the host immune response by promoting proinflammatory or anti-inflammatory processes (19). To explore the role TgLOX1 plays during animal infection, we examined TgLOX1-HA localization within infected leukocytes. We infected mice with the C- and N-terminal HA-tagged parasites and at 3 days postinfection, we collected the infected peritoneal leukocytes. Within infected peritoneal leukocytes, the TgLOX1-HA protein is no longer localized within the parasite, but instead is found within vesicle-like structures within the cytoplasm of leukocytes in both complemented strains (Fig. 6A, B and Movie 3). In extracellular parasites isolated from the peritoneal cavity, TgLOX1 is localized on the surface of the parasite (Fig. 6C). Of note, no staining of LOX1 was detected in uninfected leukocytes in the same samples (data not shown).

**Figure 6.**
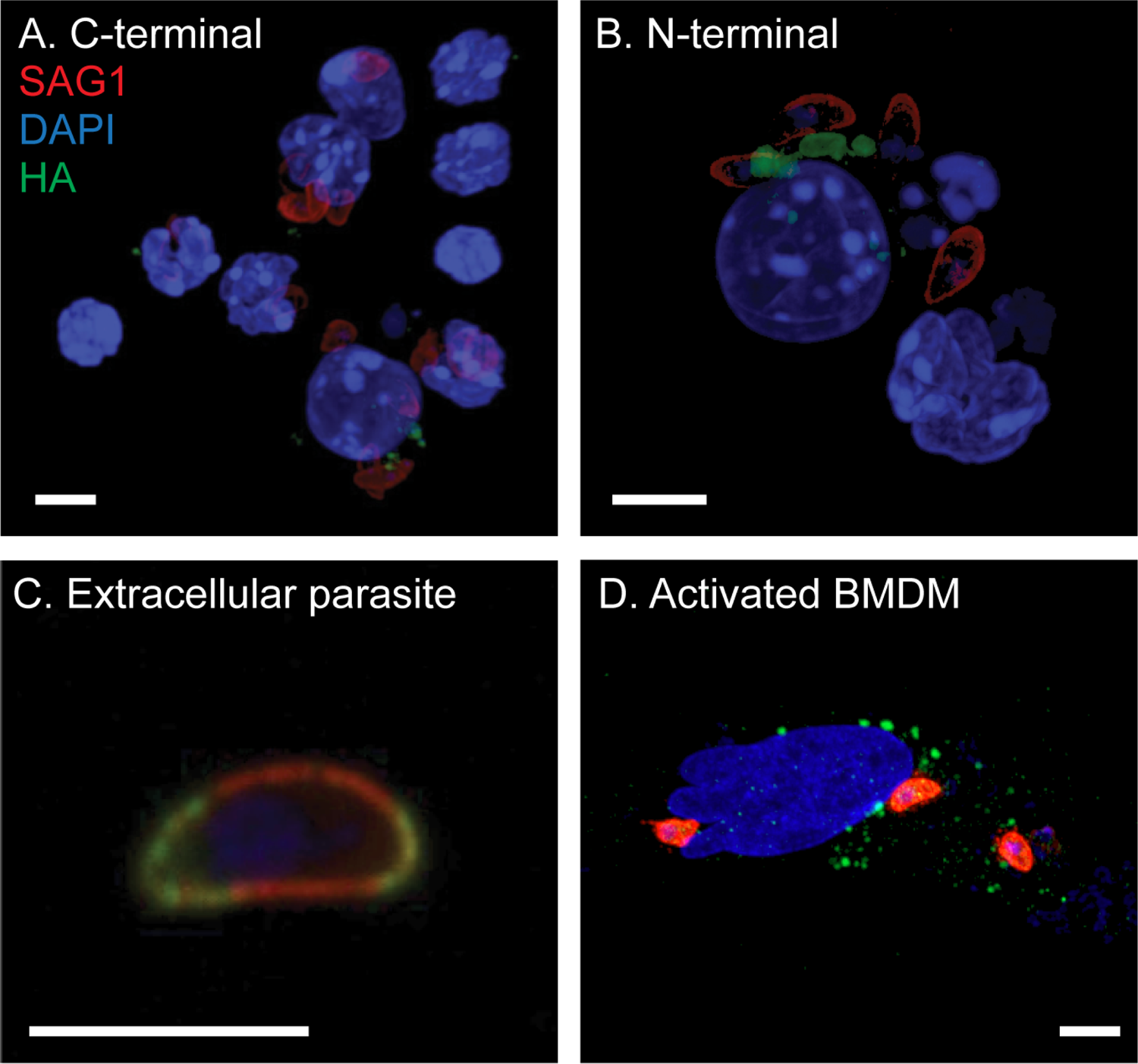
TgLOX1 is released by the parasite inside infected leukocytes. 3D projections of leukocytes isolated 3 days postinfection with 2×10^6^ C-terminal HA-tag (A) or N-terminal HA-tag parasites (B) from the peritoneal cavity of infected mice. (C) Example extracellular parasite isolated from the peritoneal cavity of infected mice with TgLOX1 with an N-terminal HA-tag (green) distributed on the parasite membrane partially co-localizing with SAG1 (red). (D) Shown a representative infected then activated BMDM in cell culture. BMDMs were infected with TgLOX1 with an N-terminal HA-tag for 3 hours, then stimulated by LPS and IFNγ and imaged after 48 hours. All scale bars are equal to 5 μm.

The fact that TgLOX1 was not detected within infected HFF but within vesicle-like structures within the cytoplasm of leukocytes isolated from mice prompted us to examine if we could recapitulate this localization change in tissue culture immune cells. We isolated bone marrow-derived macrophages (BMDMs), infected them with HA-tagged parasites, and then stimulate them with LPS and IFNγ. After 48 hours of stimulation, it was possible to detect TgLOX1 inside BMDMs in vesicle-like structures (Fig. 6D). These results were consistent with what we saw in immune cells isolated from mice (Fig. 6A and B). The fact that TgLOX1 is localized in vesicles within the leukocytes was interesting especially because TgLOX1 is not predicted to contain a signal peptide. Perhaps this localization change is important for parasite survival and/or manipulation of the host immune response.

## Discussion

Our laboratory has become interested in the lipoxygenases of *T. gondii* because of their potential role in modifying linoleic acid into lipid mediators that trigger sexual development (13). Our bioinformatic search for potential enzymes involved in processing linoleic acid led us to TgLOX1, a protein with sequence features like those found in lipoxygenases. In addition to the structural motifs, RNASeq data showed that this gene was highly transcribed in the sexual stages of *T. gondii* in contrast to the tachyzoite stage. Tissue culture characterization of ΔTgLOX1 did not reveal any phenotypes in fibroblast cells, as we had expected from a protein that was negligibly transcribed in the tachyzoite stage. However, the ΔTgLOX1 did have a deleterious phenotype during animal infection. When C57BL/6 mice were infected with ΔTgLOX1, the mice were able to clear the infection by 7 days postinfection. This phenotype prompted us to find the localization of the protein within the parasite. We found TgLOX1 to be present in the cytoplasm of parasites *in vitro*, but this changed once the parasite was injected via i.p. into C57BL/6-WT mice. *In vivo*, TgLOX1 was present in the membrane of extracellular parasites, and more interestingly, the protein appeared to be within vesicles of the infected host immune cell.

Considering our observations, we wondered how a non-essential protein for the parasite could present such a strong phenotype *in vivo*. We believe the answer is in the fundamental biology of the parasite’s infection mechanisms. When *T. gondii* invades, it uses three particular organelles in the apical complex in a coordinated fashion, micronemes, rhoptries, and dense granules (20). Microneme proteins help in the initial attachment and trigger proteins in the rhoptries to be released to help the parasite avoid detection by the immune system. Once inside the cell, proteins from the dense granules contribute to the hijacking of the host cell and pave the way for the parasite to start replication (20, 21). In the host cell, IFNγ pathways are known as the first line of defense against *T. gondii*, and the most studied to date is the JAK-STAT pathway (4, 22). ROP16 has been shown to interfere with STAT1 transcripts, altering the regulation of genes related to IFNγ and IL-12 *in vitro* (23, 24). There are three main ways in which IFNγ limits parasite replication. First by Induction of Reactive Oxygen Species and Reactive Nitrogen Species to combat intracellular parasites (25, 26). Second, induction of Indoleamine 2,3-dioxygenases (IDO1 and IDO2) to degrade tryptophan and restrict nutrients used by the parasite (27–30). Third, activation of Immunity Related GTPases (IRGs) that disrupt the Parasitophorous Vacuole Membrane (PVM) and clear the parasite from the cell (31, 32). Thus, disrupting IFNγ mediated pathways is crucial for parasite survival.

From the protein effectors that *T. gondii* uses to evade the immune response, we will focus on ROP and GRA proteins given their similarities to the behavior of TgLOX1 both *in vitro* and *in vivo*. In mice, the first line of attack against IFNγ mediated signals comes from ROP 5, 17, and 18 working in synergy to block IRGs (32, 33). Later in the infection, a GRA protein (TgIST) forms a high molecular weight complex inhibiting the STAT1 pathway that helps clear the parasite (24, 34, 35). Both classes of proteins seem to be non-essential *in vitro* but are important *in vivo* due to the interplay between them and the host’s immune response (36–38). This behavior matches what we observe in TgLOX1, of particular note is ROP38 which is highly upregulated in bradyzoites yet is still deemed non-essential (37, 39).

In our experiments, wild-type mice infected with 1×10^6^ ΔTgLOX1 parasites started to clear the infection after just 3 days. This rapid clearance resembles what Olias et al. observed when the absence of TgIST led to enhanced clearance and reduced growth of type II parasites (34). This clearance occurred in cells that were infected and then activated with IFNγ but not in naïve cells that had been prestimulated. These results indicate that TgIST acts prior to IFNγ signaling by blocking the transcriptional process. We found that there was no clearance of ΔTgLOX1 in IFNγKO mice. This result suggests that the mechanism that TgLOX1 uses is somehow altering IFNγ signaling resulting in uncontrolled parasite replication leading to the host’s demise (40, 41).

Previous work in our lab highlighted the potential importance of lipid metabolism in manipulating the host immune response by examining a patatin-like protein that protected *T. gondii* from degradation in activated macrophages (18). The absence of the protein, named TgPL1, was correlated with failure to suppress nitric oxide production and degradation in activated macrophages (42). After 33 hours in activated macrophages, parasites lacking TgPL1 showed extensive vesiculation and breakdown of the PVM. This patatin-like protein had a punctate cytoplasmic localization but was distinct from acidocalcisomes, micronemes, and rhoptries suggesting that its function is independent of secretory events associated with parasite invasion. However, TgPL1 changes localization from within the parasite to the PV and PVM after macrophage activation, colocalizing with GRA4 (42). Analysis of ΔTgPL1 parasites in mice showed no phenotype during acute infection, but an altered cytokine response during long-term chronic infection and a localization change of TgPL1 to the cyst wall (43). Unlike TgPL1, the absence of TgLOX1 results in parasite clearance during acute infection but the mechanism is still unknown. Our current hypothesis is that TgLOX1 helps *T. gondii* avoid detection by the host immune system either by scrambling the composition of available oxylipins within the host cell or by affecting the JAK-STAT signaling pathway (34). To our knowledge, this is the first report of a *T. gondii* lipoxygenase and its importance *in vivo*. There is still much to be discovered in the association between the parasite’s lipid metabolism and its role in immune evasion.

## Materials and Methods

### Ethics Statement

Mice were treated in compliance with the guidelines set by the Institutional Animal Care and Use Committee (IACUC) of the University of Wisconsin School of Medicine and Public Health (protocol #M005217). The institution adheres to the regulations and guidelines set by the National Research Council.

### Parasite and host cell culture

*T. gondii* tachyzoites were cultured in human foreskin fibroblasts (HFF) in Dulbecco’s modified Eagle’s medium (DMEM, Gibco) supplemented with 10% fetal bovine serum, 2mM L-glutamine, and 1% penicillin-streptomycin at 37°C with 5% CO_2_. To avoid the appearance of any tissue culture specific characteristics, low passage parasites were frozen in 10% DMSO, 20% FBS, and 70% DMEM and used until passage 15.

### Sequence alignment and candidate gene selection

To select the potential LOX candidates involved in the sexual development of *T. gondii* we performed a database search. Annotated protein data from the *T. gondii* TGME49 strain was downloaded from ToxoDB.org and the amino acid sequence of each protein was scanned for the presence of a C-terminal Isoleucine or Valine. The resulting protein IDs from this scan were then used in a search strategy on ToxoDB. The gene ID was cross referenced with transcripts that were more than 100-fold upregulated in the sexual stage compared to the tachyzoite stage using the feline enterocyte, tachyzoite, bradyzoite stage transcriptome (16). Then the candidates were organized from highest to lowest TPM value and our candidate gene was selected.

### Generation of TgLOX1 knockout and complement strains

A pBC_TubCAT_HXGPRT empty vector was digested with SpeI and KpnI to insert the flanking regions of our gene of interest on either side of the chloramphenicol acetyl transferase (CAT) positive selectable marker, with hypoxanthine-xanthine-guanine phosphoribosyl transferase (HXGPRT) as the negative selectable marker. 4kb genomic fragments 5’ and 3’ of the TgLOX1 locus were amplified by PCR with Phusion polymerase (Thermo Scientific) using primers LOX1 5’ flank F/R and LOX1 3’ flank F/R. The resulting pBC_TubCAT_HXGPRT_315970 vector was linearized prior to electroporation of parental PruΔHPT:luciferase parasites (42) with the BTX ECM 630 Electroporation System. Successful knockout strains were selected with chloramphenicol and 6-thioxanthine and clonal populations were isolated by limiting dilution. Loss of the full TgLOX1 gene was confirmed by PCR using LOX1 KO F/R primers (Fig. S1). Complement strains were created by amplifying cDNA from the TgLOX1 coding region using the LOX1 ORF F/R primers and inserting the coding region into a pBC_SK vector containing a tubulin promoter and SAG1 3’ UTR with a DHFR positive selectable marker. The resulting pBC_SK_DHFR_315970 vector was linearized prior to electroporation with the BTX ECM 630 Electroporation System.

Successful complement strains were selected for non-homologous recombination with pyrimethamine, and clonal populations were isolated by limiting dilution. The presence of the TgLOX1 coding region was confirmed by PCR using LOX1 KO F/R primers. All primers used in this work can be found in Table S1.

### Plaque Assay

Plaque assays were performed by seeding HFF cells in 12-well plates until a confluent monolayer was formed. Cells were infected with 500 parasites of parental PruΔHPT:luciferase, ΔTgLOX1, or complement untagged parasite strains, and left undisturbed for 7 days for the plaques to form. The medium was removed, and the cells were fixed with methanol and stained with crystal violet for 20 minutes. Then the samples were rinsed with water and left to air dry overnight and then photographed. Plaques were counted using ImageJ software by analyzing the number of individual plaques formed.

### Replication Assay

Replication assays were performed by seeding HFF cells on sterile glass coverslips in 24-well plates and infecting the monolayer with 10^5^ parasites of parental PruΔHPT:luciferase, ΔTgLOX1, or complement untagged parasite strains in triplicate. After 2 hours of incubation, each well was washed with PBS to remove extracellular parasites, and then new media was added. After 12, 24, or 36 hours postinfection the samples were fixed and stained with 1:500 rabbit anti-SAG1 primary antibody and 1:1000 goat anti-rabbit Alexa Fluor 488 and counterstained with DAPI. After mounting, the slides were blinded and the number of parasites per vacuole was counted. The statistics were performed using Analysis of Variance (ANOVA) in the GraphPad prism software.

### Mouse Infections and parasitemia measurements

Mice from 8 to 12 weeks old were used for all the experiments and to the best of our abilities maintained the same proportion of male and female mice to account for sex-dependent variability. For the infections, parasites were scraped and syringed from HFF monolayers and counted for i.p. injection of parental PruΔHPT:luciferase, ΔTgLOX1, or complement untagged parasite strains. The number of parasites used for infection varied depending on the experiment being performed and the strain of mice used (WT vs. IFNγKO).

#### Survival curves

Between 4-5 mice were used for each parasite strain and the experiment was repeated at least twice. NMRI and C57BL/6 Wild-type (WT) mice were inoculated via i.p. injection with 1×10^5^ parasites. IFNγKO mice were inoculated with 1×10^2^-1×10^4^ parasites due to their higher susceptibility to infection. Mice were monitored daily up to 28 days postinfection for disease symptoms scored on a 1-4 scale, with 1 indicating no pain and distress and 4 indicating moribund condition (44). Surviving mice were checked for parasitemia. Prism GraphPad software was used to analyze survival curves by log-rank (Mantel-Cox).

#### Measuring acute parasitemia by IVIS

Between 4-5 mice were used for each parasite strain and the experiment was repeated at least twice. WT mice were inoculated by i.p. injection with 1×10^5^ or 1×10^6^ parasites and IFNγKO mice were inoculated with 1×10^4^ parasites. At 7 days postinfection parasitemia in mice was detected by bioluminescence using IVIS (PerkinElmer). Wild-type mice infected with the higher parasite dose of 1×10^6^ parasites were imaged at 3 days postinfection. Prior to imaging, mice were injected i.p. with 200 μL of D-Luciferin potassium salt (15.4 mg/mL in PBS) and anesthetized with 4% isoflurane. Anesthesia was maintained throughout the imaging process at 2% isoflurane. After 15 minutes images were collected, and the mice were returned to their cages to recover. A background measurement was determined by injecting an uninfected mouse with D-Luciferin and collecting the images as previously described. Data collection was performed by drawing a region of interest (ROI) box for each mouse and recording the total number of photons per second (Total Flux) (45).

#### Chronic infection parasitemia by luciferase and DBA

Experiments were performed using NMRI and Swiss Webster mice inoculated via i.p. injection with 1×10^4^ parasites. After 28 days, brains were collected and homogenized in 1 mL of ice-cold PBS. For the luciferase assay, a 750 μL aliquot from the sample was transferred to a clean Eppendorf tube and mixed with 250 μL of 5x lysis buffer (125mM Tris, 10mM DTT, 50% Glycerol, 5% Triton-X100) and incubated for 20 minutes in ice. The sample was mixed, separated into two 500 μL aliquots, and left on ice. A stock of 2x luciferase reaction buffer (10 mM MgCl_2_, 0.3 mM ATP, 200 mM, Luciferin 1.5 mg/mL final concentration) was prepared, left at room temperature, and protected from light until samples were ready to be measured. Each sample aliquot was mixed with 500 μL of the 2x luciferase reaction buffer, mixed by inversion and its luminescence was measured on a GloMax® 20/20 Luminometer (Promega). For the DBA assay, the samples were fixed in 3.7% formaldehyde in PBS for 20 minutes and permeabilized and blocked with 0.2% v/v Triton x-100 (Sigma) and 3% BSA in PBS at room temperature for one hour. The samples were incubated with a 1:500 dilution of biotinylated DBA (Vector Laboratories) followed by a 1:500 dilution of streptavidin Alexa Fluor 594 (Thermofisher) for one hour each, washing after each incubation.

### Cytokine Bead Assay

Serum cytokine levels were measured using the BD cytometric bead array mouse inflammation kit (BD Biosciences). Blood serum was collected from the mice used in the IVIS experiment on days 0, 3, and 7 postinfection. Samples were processed according to the manufacturer’s instructions and analyzed using an Attune flow cytometer (Thermo-Fisher) at the University of Wisconsin Carbone Cancer Center. Further analysis was performed using the FlowJo software. The statistics were performed using one-way ANOVA in the GraphPad prism software.

### Western Blot

HFF monolayers were infected and after 24 hours postinfection cells were scraped and lysed by syringe (28G needle). Detached cells were then centrifuged 500 x g for 10 min and the pellet was solubilized using lysis buffer (2% 2-mercaptoethanol, 1% SDS, 20 mM EGTA, 2 mM Tris–HCl at pH 7.5, 0.1 mM PMSF, 0.1 mM TPCK, and 0.1 mM TLCK) and sonicated in ice for 15 s at 40 Hz (Branson Sonifier 250). Approximately 20 µg of protein were loaded into a gel (SDS-PAGE), transferred to a nitrocellulose membrane, and blocked with 6% skim milk dissolved in TBS-T (10 mM Tris–HCl, 75 mM NaCl and 0.1% Tween 20 at pH 8.0,) for 1 hour. Primary antibody against-SAG1 (rabbit, 1:3000 kindly donated by John Boothroyd), -HA Tag (mouse monoclonal, Enzo Life Sciences, NY) was incubated overnight at 4°C and then thoroughly washed with TBS-T. Secondary antibodies conjugated to horseradish peroxidase (HRP, rabbit or mouse 1:7000, Thermo Scientific) were incubated for 2 hours at room temperature.

Antibody 6xHisTag was already conjugated to HRP (Goat polyclonal, 1:2000, Bethyl Laboratories). Chemiluminescence (ECL. GE, Buckingham-shire, UK) was detected using the Odyssey XK instrument (LI-COR Imaging system).

### Indirect immunofluorescence assays

To obtain the samples: (a) HFF monolayers were seeded in 24 well-plates and infected as described above, or (b) C57BL/6 mice were infected via i.p. injection with 2 x 10^6^ parasites and after 3 days, infected leukocytes were isolated by peritoneal lavage. Cells were fixed with 4% paraformaldehyde (PFA), permeabilized with 0.5% Triton X-100, and blocked with 5% BSA. Cells were incubated with the primary antibodies against-SAG1 (rabbit, 1:300), -MIC2 (rabbit, 1:300) or HA-Tag (mouse monoclonal, 1:1000) overnight at 4°C and then thoroughly washed with PBS. Secondary antibodies against -Alexa Fluor 488 (Rabbit anti-mouse, 1:1000, Thermo Scientific) and -Alexa Fluor 594 (Goat anti-rabbit, 1:1000, Thermo Scientific) were incubated for 2 hours at room temperature. Leukocytes were seeded on poly-L-lysine coated coverslips. Cells were counterstained with DAPI and mounted using VECTASHIELD antifade Mounting Medium (Vector Laboratories Inc., CA). Cells were imaged using an epifluorescence microscope (Imager.M2, Carl Zeiss, DEU) or a confocal microscope (ZeissLSM 800 Laser Scanning Microscope). 3D projections were reconstructed from z-stack sections using the Zen imaging software (Carl Zeiss).

### Infection and imaging of BMDMs

BMDMs were harvested from 8- to 10-week-old mice and grown in 20% L929 conditioned RPMI medium as described previously (46). A total of 1×10^5^ viable BMDM were seeded on poly-L-lysine coated coverslips and infected with 1 × 10^5^ N-terminal HA-tag tachyzoites. After 3 h postinfection, cells were stimulated with 100 ng/ml lipopolysaccharide (LPS) and 100 U/ml IFN-γ. At 48 h postinfection, cells were fixed, processed, and imaged as above described.

## Acknowledgments

We sincerely thank John Boothroyd and David Sibley for antibodies. This research was supported by the National Institutes of Health (NIH) National Research Service Award T32 AI007414 (SKW), and R01AI144016-01(LJK), SciMed Graduate Research Scholars Fellowship UW-Madison (CMC), and Foster Wisconsin Distinguished Fellowship from the Food and Research Institute (CMC).

---

**Figure S1. Generation of the knockout strain ΔTgLOX1.** The homology region for the knockout was designed to flank the upstream and downstream regions of the LOX1 gene. Amplification of the LOX1 gene in wild type (parental) strain showed a 3361 bp band in contrast to the 2879 bp in the ΔTgLOX1, indicating the loss of the gene.

**Figure S2. TgLOX1 is not essential for growth in tissue culture.** (A) Triplicate monolayers of HFFs were infected with *T. gondii*. At 12, 18, or 36 hours postinfection, tachyzoites per vacuole were scored on blinded slides for at least 100 random vacuoles per replicate. Average tachyzoites per vacuole are shown at each time point. (B) To see the formation of plaques, HFF monolayers were infected with 500 parasites of each strain. After 7 days postinfection, cells were stained with crystal violet to reveal *T. gondii* plaque formation. (C) Shown the number of plaques formed and quantified using ImageJ software. The statistics were performed using one way ANOVA in the GraphPad prism software. The *p* value was considered as follows: * <0.05 and ** <0.005.

**Figure S3. ΔTgLOX1 parasites have reduced parasitemia during chronic infection.** (A) NMRI mice were infected with 1×10^4^ parasites of each strain and at 28 days postinfection, the brains were removed for luciferase assays. ** <0.005. (B) Survival curve of NMRI mice infected with 1×10^5^ parasites of each strain. Mice infected with parental and complement parasites became moribund between days 9-15. (C) A few cysts were found in the brains of ΔTgLOX1-infected mice at 28 days postinfection. Brains were stained for DBA (red) and imaged with Differential Interference Contrast (DIC). Scale bar equals 5 μm. (D) Shown are Swiss Webster mice infected with 1×10^4^ parasites of each strain and at 28 days postinfection, the brains were removed for luciferase assays. * <0.05.

**Figure S4. ΔTgLOX1 parasitemia is reduced even with large inoculums.** Shown two independent experiments of 3-5 C57BL/6-WT mice, either all males or all females, with a total of 7-8 mice per strain. Mice were i.p. infected with 1×10^6^ of luciferase-expressing parasites of each strain and 3 days postinfection they were imaged ventrally by IVIS. (A and B) Shown are the images for each strain and separately for gender. For all mice, the abdominal hair was removed to avoid signal interference and the exposure time was the same. (C) Shown the total flux was obtained by measuring the luminescence intensity in the peritoneal cavity of mice. The statistics were performed using one way ANOVA in the GraphPad prism software. The *p* value was considered as ** <0.005.

**Figure S5. MCP-1 and IL-6 are upregulated in the serum of IFNγKO mice during acute infection.** Graphs show a comparative profile of the cytokine response in wildtype and IFNγKO mice. Blood serum of female and male C57BL/6-WT and IFNγKO mice were collected to analyze the cytokine response. Mice were infected with 1×10^4^ parasites of each strain and analyzed at 7 days postinfection. MCP-1 and IL-6 are more abundant in IFNγKO mice, but there are no significant differences between strains in IFNγKO mice. TNF-α is significantly lower in ΔTgLOX1-infected mice compared to parental- and complement-infected WT mice, but TNF-α is significantly higher in ΔTgLOX1-infected mice compared to parental- and complement-infected IFNγKO mice. The statistics were performed using one way ANOVA in the GraphPad prism software. The *p* value was considered as follows: ** <0.005 and *** <0.0005.

